# Elevated Na^+^/K^+^ Ratio in Alzheimer’s Disease: A Potential Biomarker for Braak Stage

**DOI:** 10.1101/2025.11.04.686539

**Authors:** Yuma Mizuno, Shiyue Pan, Tong Zhou, Patrick G. Kehoe, Yumei Feng Earley

**Affiliations:** Department of Physiology and Cell Biology, University of Nevada, Reno, NV, USA; Department of Medicine, University of Rochester Medical Center, Rochester, NY, USA; Department of Pharmacology & Physiology, University of Rochester Medical Center, Rochester, NY, USA; Cerebrovascular and Dementia Research Group, Translational Health Sciences, Bristol Medical School, University of Bristol, Bristol, UK

**Keywords:** cerebrospinal fluid sodium, brain sodium, Alzheimer’s Disease, Na^+^/K^+^ ratio

## Abstract

Alzheimer’s disease (AD) is a neurodegenerative disorder characterized by cognitive decline, synaptic dysfunction, and the accumulation of amyloid plaques and neurofibrillary tangles. While prior research has focused mainly on protein aggregation and neuroinflammation, emerging evidence suggests that ionic imbalances, particularly involving sodium (Na+) and potassium (K+), may contribute to AD progression. Na+ and K+ are critical for maintaining neuronal membrane potential, regulating action potential firing, and supporting neurotransmitter function. Although studies primarily focused on absolute Na+ concentrations, the Na+/K+ ratio may provide a more sensitive marker of ionic dysregulation. Given that the Na+/K+ gradient is actively maintained by the Na/K-ATPase pump, a target known to be vulnerable in AD, we hypothesized that the Na+/K+ ratio is altered in AD. We analyzed postmortem tissue from the prefrontal cortex, thalamus, and cerebrospinal fluid (CSF) of 97 human subjects (67 AD, 30 controls). AD cases exhibited a significant increase in the Na+/K+ ratio in the thalamus and CSF, driven primarily by elevated Na+ levels. The Na+/K+ ratio positively correlated with Braak tangle stage, suggesting an association with AD progression. These findings provide novel insights into ionic dysregulation in AD and suggest that the CSF Na+/K+ ratio may serve as a valuable biomarker of disease severity and progression. Future research should explore the potential of targeting ionic homeostasis as a therapeutic strategy in AD.

## 1. Introduction

Alzheimer’s disease (AD) is a progressive neurodegenerative disorder characterized by cognitive decline, synaptic dysfunction, and the accumulation of amyloidβ (Aβ) plaques and neurofibrillary tangles [1,2]. While significant research has focused on Aβ aggregation, tau pathology, and neuroinflammation in AD [3,4], accumulating evidence suggests that dysregulation of ion homeostasis, particularly involving sodium (Na^+^) and potassium (K^+^), may be an important but underexplored contributor to disease progression [5-7].

Na^+^ and K^+^ ions are critical for maintaining neuronal membrane potential, supporting action potential generation, regulating synaptic transmission, and sustaining metabolic homeostasis [8-14]. The steep transmembrane gradients of Na^+^ and K^+^ are actively maintained by the Na^+^/K^+^-ATPase pump, a high-energy-demanding enzyme known to be vulnerable to oxidative stress and metabolic dysfunction, which are prevalent in AD brains [15-19]. Even subtle perturbations in Na^+^ and K^+^ levels can lead to impaired neuronal excitability, calcium overload, and ultimately cell death [20-25]. Previous studies have reported elevations in Na^+^ levels in the post-mortem brains of AD patients [26] that correlate with Braak tangle stage, a widely used neuropathological marker of disease severity. However, prior research has largely focused on absolute Na^+^ concentration [Na^+^], without considering the Na^+^/K^+^ ratio.

In this study, we investigated [Na^+^], [K^+^], and Na^+^/K^+^ ratio in homogenates made from postmortem brain tissues (prefrontal cortex and thalamus) and cerebrospinal fluid (CSF) from people who died with a diagnosis of AD or non-dementia. Our findings reveal a significant increase in the Na^+^/K^+^ ratio in AD brains and CSF compared to non-dementia controls, suggesting dysregulation of ion balance in AD pathophysiology. Furthermore, our study uniquely highlights an association between impairments in Na^+^/K^+^-ATPase and alterations in Na^+^/K^+^ ratio in AD, shedding new light on mechanisms that may underlie neurodegeneration. Given the emerging recognition of the Na^+^/K^+^ ratio as a superior biomarker in other conditions, such as hypertension [27-29], these findings may stimulate further studies focused on ionic imbalance in AD risk assessment and progression.

## 2. Results

### 2.1. Study subject characteristics

All study subject characteristics are detailed in Table 1, which includes a summary of sex, race, age, and Braak tangle state. All patients were clinically diagnosed with AD. These were confirmed by neuropathological assessment post-mortem for the presence (Patients) or absence (Controls) of AD-related pathological abnormalities. Control and AD subjects were similar with respect to sex (*P* = 1), race (*P* = 0.369), whereas the mean age of subjects (related to age at death) was significantly younger in the AD (70±10 years) compared with the Control (84±8) group (P < 0.001).

**Table 1.** Characteristics of study subjects.

|  | Control | AD |
| --- | --- | --- |
| **Frontal Cortex, Thalamus, CSF** |  |  |
| Number of subjects/groups | 30 | 67 |
| **Demographics:** |  |  |
| sex, men/women | 15/15 | 35/32 |
| race, White (European)/unknown | 27/3 | 64/3 |
| Age, years | 84±8 | **77±10\*** |

| Braak Tangle Stage | 0-III | IV-VI |
| --- | --- | --- |
| Values are means ± SEM where appropriate. *$P < 0.001$ AD vs. Control. | | |

### 2.2. Increased [Na^+^] and Na^+^/K^+^ ratio in CSF and brain tissues of AD patients

The Na^+^/K^+^ ratio is commonly used as an indicator of cardiovascular disease risk, including hypertension [27,30]. Accordingly, we assessed the levels of Na^+^ and K^+^ in the prefrontal cortex, thalamus, and CSF of AD patients (n = 67) and controls (n = 30). [Na^+^] in the prefrontal cortex (Figure 1A-C) was similar between AD patients and controls (*P =* 0.33), whereas [K^+^] trended lower (*P* = 0.06) in AD patients compared with control subjects. Interestingly, despite the absence of a significant difference in either [Na^+^] or [K^+^], the Na^+^/K^+^ ratio in the prefrontal cortex was significantly higher in AD subjects (Figure 1C; *P* = 0.003). In thalamus tissue (Figure 1D-F), [Na^+^] was significantly higher in AD subjects compared with controls (*P* = 0.046), but [K^+^] was similar between the two groups, resulting in a significantly higher Na^+^/K^+^ ratio in AD subjects than in controls (Figure 1F; *P* = 0.032). In the CSF (Figure 1G-I), [Na^+^] was markedly higher (*P* < 0.0001) and [K^+^] was markedly lower (*P* = 0.0006) in AD subjects; consequently, the Na^+^/K^+^ ratio was significantly higher in AD patients compared with control subjects (Figure 1I, *P* < 0.0001).

**Figure 1.**
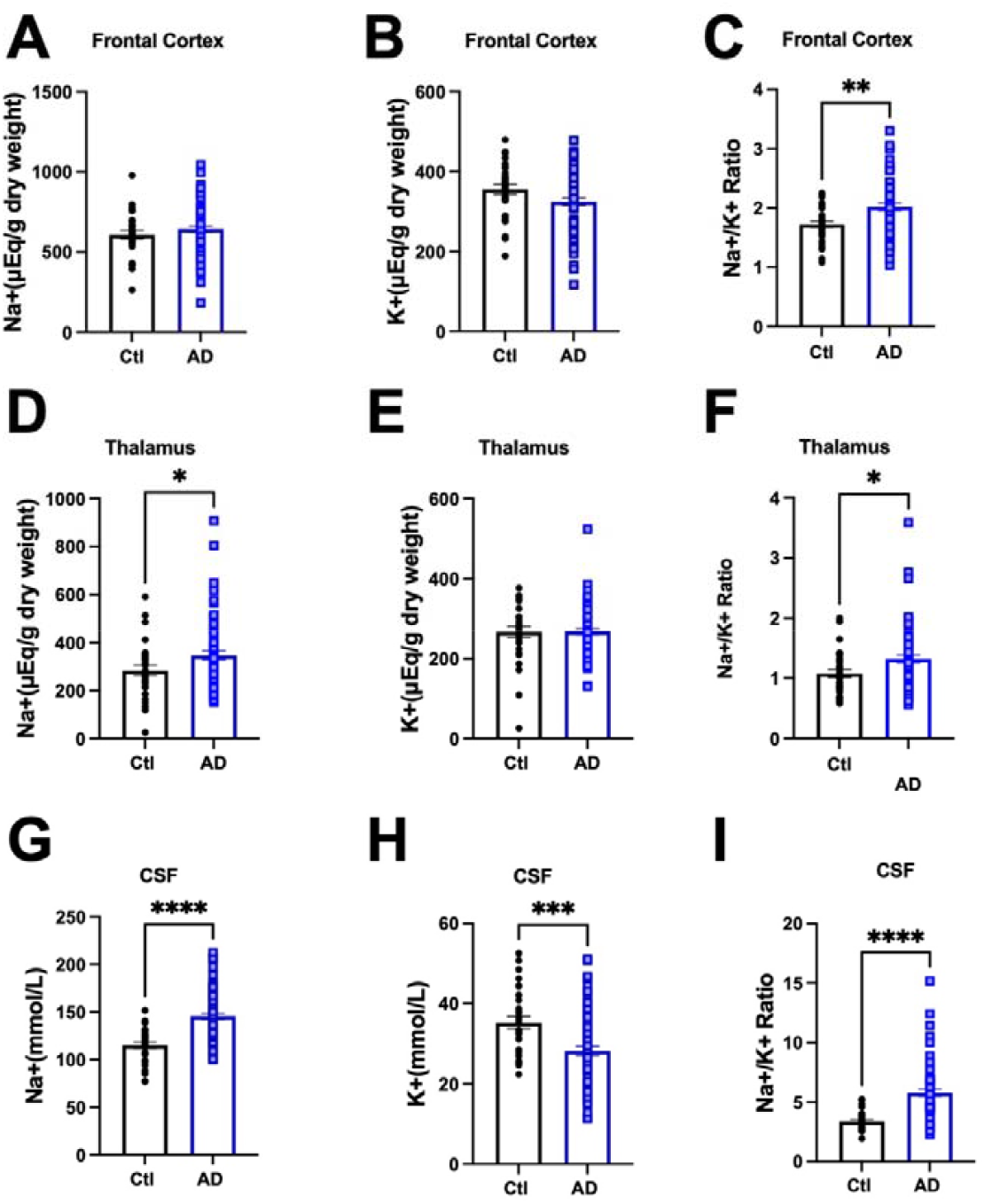
Na^+^, K^+^, and Na^+^/K^+^ ratio in AD humans’ prefrontal cortex, thalamus, and CSF. **(A-C)** Na^+^ (A), K^+^ (B), and Na^+^/K^+^ ratio (C) in the frontal cortex of control (Ctl) versus AD humans. **(D-F)** Na^+^ (D), K^+^ (E), and Na^+^/K^+^ ratio (F) in the thalamus of control (Ctl) versus AD humans. **(G-I)** Na^+^ (G), K^+^ (H), and Na^+^/K^+^ ratio (I) in the cerebrospinal fluid (CSF) of control (Ctl) versus AD humans. Ctl: N = 30; AD: N = 67. Unpaired *t*-test, \**P* < 0.05, \*\**P* < 0.01 \*\*\**P* < 0.001 \*\*\*\**P* < 0.0001 vs. Ctl.

### 2.3 Relationship between brain tissue and CSF [Na^+^] and [K^+^]

To assess whether Na^+^ and K^+^ levels in CSF reflect those in brain tissue, we performed linear regression analyses comparing CSF [Na^+^] and [K^+^] with levels of sodium and potassium in the frontal cortex and thalamus across all subjects (Figure 2). In the frontal cortex (Figure 2A,B), CSF [Na^+^] showed no correlation with tissue Na], whereas CSF [K^+^] was positively correlated with cortex tissue K^+^ (P = 0.0099, R^2^ = 0.07). In the thalamus (Figure 2C,D), CSF [Na^+^] was not correlated with tissue Na^+^, while CSF [K^+^] was negatively correlated with tissue K^+^ (P = 0.033, R^2^ = 0.05). Although the correlation between CSF [K^+^] and brain tissue K^+^ (Figure 2B,D) reached statistical significance, the R^2^ values were very low, suggesting an extremely weak and no meaningful correlation between the CSF [Na^+^] and [K^+^] and the respective tissue Na^+^ and K^+^ levels. To evaluate whether these relationships were influenced by AD, we repeated the analyses within the AD and control subgroups; no significant correlations were observed in either the cortex or the thalamus (Supplemental Figures 1 and 2).

**Figure 2.**
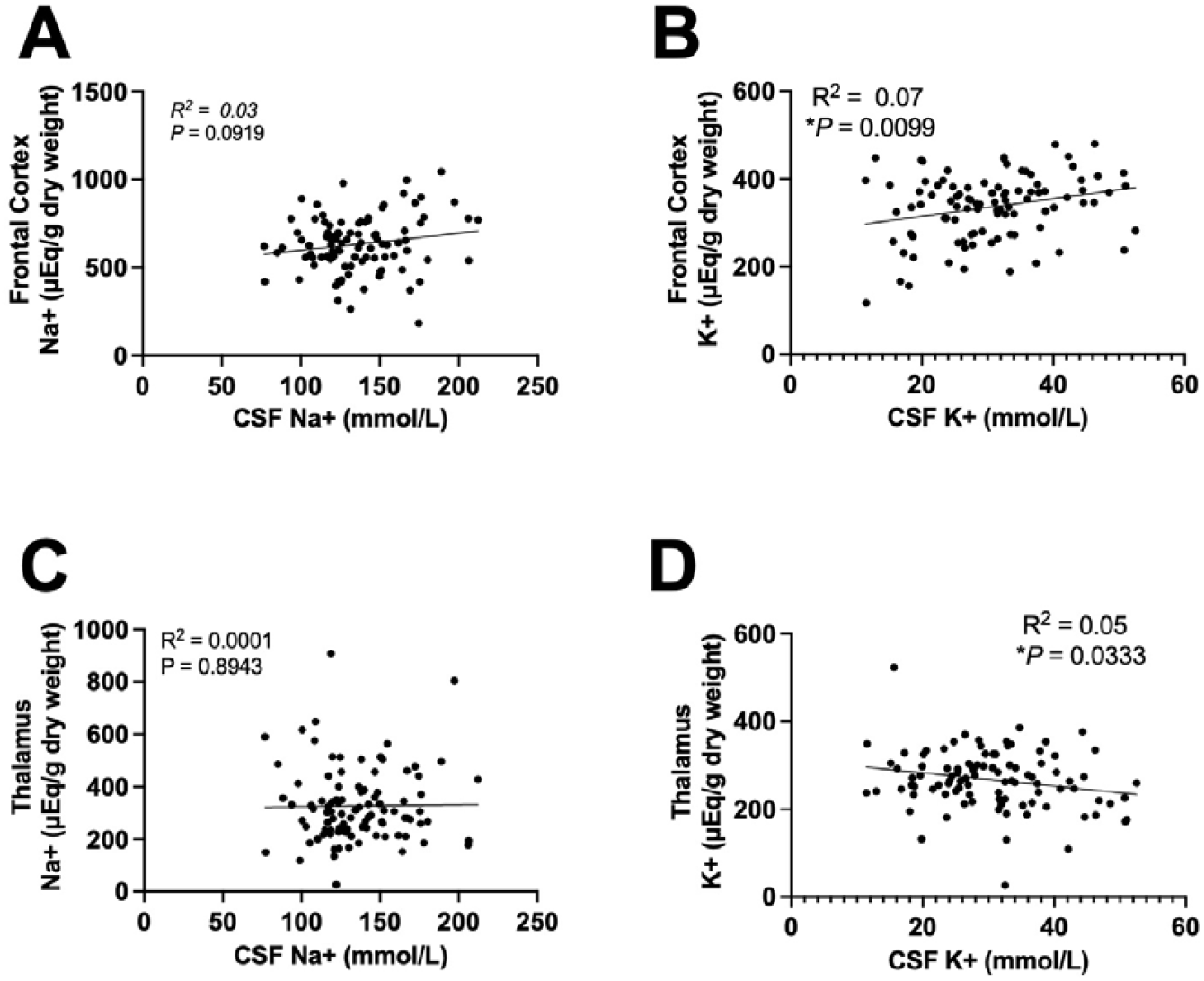
Correlations between tissue and CSF Na^+^ and K^+^ concentrations in control and AD humans. **(A, B)** Correlations between Na^+^ (A) and K^+^ (B) levels in the frontal cortex and CSF. **(C, D)** Correlations between Na^+^ (C) and K^+^ (D) levels in the thalamus and CSF. Simple Linear Regression, N = 97. Two-tailed, and differences were considered statistically significant at *P*-values < 0.05.

### 2.4 [Na^+^] and Na^+^/K^+^ ratios are associated with the severity of Braak stage in AD patients

Braak stage is used as a basis for making a post-mortem diagnosis of AD and in research as an indicator of the relatively predictable progression of tau-based neurofibrillary tangle pathology in AD [31,32]. Age is considered the most significant non-modifiable risk factor for AD [33,34], whereas sex differences have also been reported in the incidence and severity of AD [35]. As shown in Table 1, AD subjects used in this study were younger than controls at the time of tissue collection, although we did not see noticeable sex differences in our cohort. To accurately determine correlations between [Na^+^], [K^+^], or Na^+^/K^+^ ratio with Braak stage severity (0-VI), we performed multiple regression analyses, controlling for sex and age. Across the frontal cortex and thalamus, and within CSF, [Na^+^] and Na^+^/K^+^ ratios were consistently and positively correlated with Braak stage (Table 2). Neither frontal cortex nor thalamus [K^+^] was correlated with Braak stage, whereas [K^+^] in the CSF remained negatively correlated with Braak stage. [Na^+^], [K^+^], and Na^+^/K^+^ ratios in the different Braak stages (0-VI), visualized by plotting as bar graphs (Figure 3), showed that Na^+^ and Na^+^/K^+^ ratios gradually increased with the progression of Braak stage from 0 to VI. Collectively, these data indicate that [Na^+^] and Na^+^/K^+^ ratio in CSF and the brain tissues examined are positively correlated with AD severity, determined by Braak stage.

**Table 2.** Multivariate linear regression between Na^+^, K^+^, or Na^+^/K^+^ ratio with Braak stage, controlling for sex and age in the frontal cortex, thalamus, and CSF.

| Region | Variable | Estimate | 95% CI | Std. Error | t value | Pr(>\|t\|) |
| --- | --- | --- | --- | --- | --- | --- |
| **Frontal cortex** |  |  |  |  |  |  |
| $Na^+$ | Intercept | 364.362 | (42.957, 685.768) | 161.852 | 2.251 | **2.67E-02\*** |
|  | Sex | -21.198 | (-84.329, 41.933) | 31.791 | -0.667 | 5.07E-01 |
|  | Age | 2.384 | (-1.069, 5.837) | 1.739 | 1.371 | 1.74E-01 |
|  | Braak tangle stage | 21.119 | (1.832, 40.407) | 9.713 | 2.174 | **3.22E-02\*** |
| $K^+$ | Intercept | 271.396 | (126.899, 415.893) | 72.765 | 3.730 | **3.29E-04\*** |
|  | Sex | -39.620 | (-68.002, -11.238) | 14.293 | -2.772 | **6.73E-03\*** |
|  | Age | 1.315 | (-0.237, 2.868) | 0.782 | 1.682 | 9.58E-02 |
|  | Braak tangle stage | -4.735 | (-13.407, 3.936) | 4.367 | -1.084 | 2.81E-01 |
| $Na^+/K^+$ ratio | Intercept | 1.592 | (0.709, 2.474) | 0.444 | 3.582 | **5.45E-04\*** |
|  | Sex | 0.157 | (-0.016, 0.331) | 0.087 | 1.803 | 7.46E-02 |
|  | Age | -0.002 | (-0.011, 0.008) | 0.005 | -0.398 | 6.92E-01 |
|  | Braak tangle stage | 0.096 | (0.043, 0.149) | 0.027 | 3.590 | **5.31E-04\*** |
| **Thalamus** |  |  |  |  |  |  |
| $Na^+$ | Intercept | 368.341 | (81.651, 655.032) | 144.370 | 2.551 | **1.24E-02\*** |
|  | Sex | -4.386 | (-60.698, 51.926) | 28.357 | -0.155 | 8.77E-01 |
|  | Age | -1.483 | (-4.563, 1.597) | 1.551 | -0.956 | 3.41E-01 |
|  | Braak tangle stage | 18.681 | (1.477, 35.885) | 8.664 | 2.156 | **3.36E-02\*** |
| $K^+$ | Intercept | 218.169 | (80.657, 355.680) | 69.247 | 3.151 | **2.19E-03\*** |
|  | Sex | 24.570 | (-2.440, 51.580) | 13.602 | 1.806 | 7.41E-02 |
|  | Age | 0.275 | (-1.202, 1.752) | 0.744 | 0.369 | 7.13E-01 |
|  | Braak tangle stage | 3.466 | (-4.786, 11.718) | 4.156 | 0.834 | 4.06E-01 |
| $Na^+/K^+$ ratio | Intercept | 1.480 | (0.460, 2.500) | 0.514 | 2.882 | **4.91E-03\*** |
|  | Sex | -0.119 | (-0.319, 0.081) | 0.101 | -1.180 | 2.41E-01 |
|  | Age | -0.005 | (-0.016, 0.006) | 0.006 | -0.973 | 3.33E-01 |
|  | Braak tangle stage | 0.061 | (-0.001, 0.122) | 0.031 | 1.964 | 5.25E-02 |
| **CSF** |  |  |  |  |  |  |
| $Na^+$ | Intercept | 75.468 | (24.327, 126.609) | 25.754 | 2.930 | **4.26E-03\*** |
|  | Sex | 0.133 | (-9.913, 10.178) | 5.059 | 0.026 | 9.79E-01 |
|  | Age | 0.316 | (-0.234, 0.865) | 0.277 | 1.141 | 2.57E-01 |
|  | Braak tangle stage | 8.378 | (5.309, 11.447) | 1.546 | 5.421 | **4.65E-07\*** |
| $K^+$ | Intercept | 41.255 | (23.133, 59.377) | 9.126 | 4.521 | **1.81E-05\*** |
|  | Sex | -2.705 | (-6.265, 0.854) | 1.793 | -1.509 | 1.35E-01 |
|  | Age | -0.001 | (-0.195, 0.194) | 0.098 | -0.006 | 9.95E-01 |
|  | Braak tangle stage | -2.224 | (-3.312, -1.137) | 0.548 | -4.061 | **1.02E-04\*** |
| $Na^+/K^+$ ratio | Intercept | 3.463 | (-0.633, 7.559) | 2.063 | 1.679 | 9.66E-02 |
|  | Sex | -0.035 | (-0.839, 0.770) | 0.405 | -0.086 | 9.32E-01 |
|  | Age | -0.014 | (-0.058, 0.030) | 0.022 | -0.649 | 5.18E-01 |
|  | Braak tangle stage | 0.649 | (0.403, 0.894) | 0.124 | 5.239 | **1.00E-06\*** |

\* $P < 0.05$ indicates that Braak tangle stage has a significant effect on the response variables (i.e., $Na^+$ , $K^+$ , or $Na^+/K^+$ ratio).

**Figure 3.**
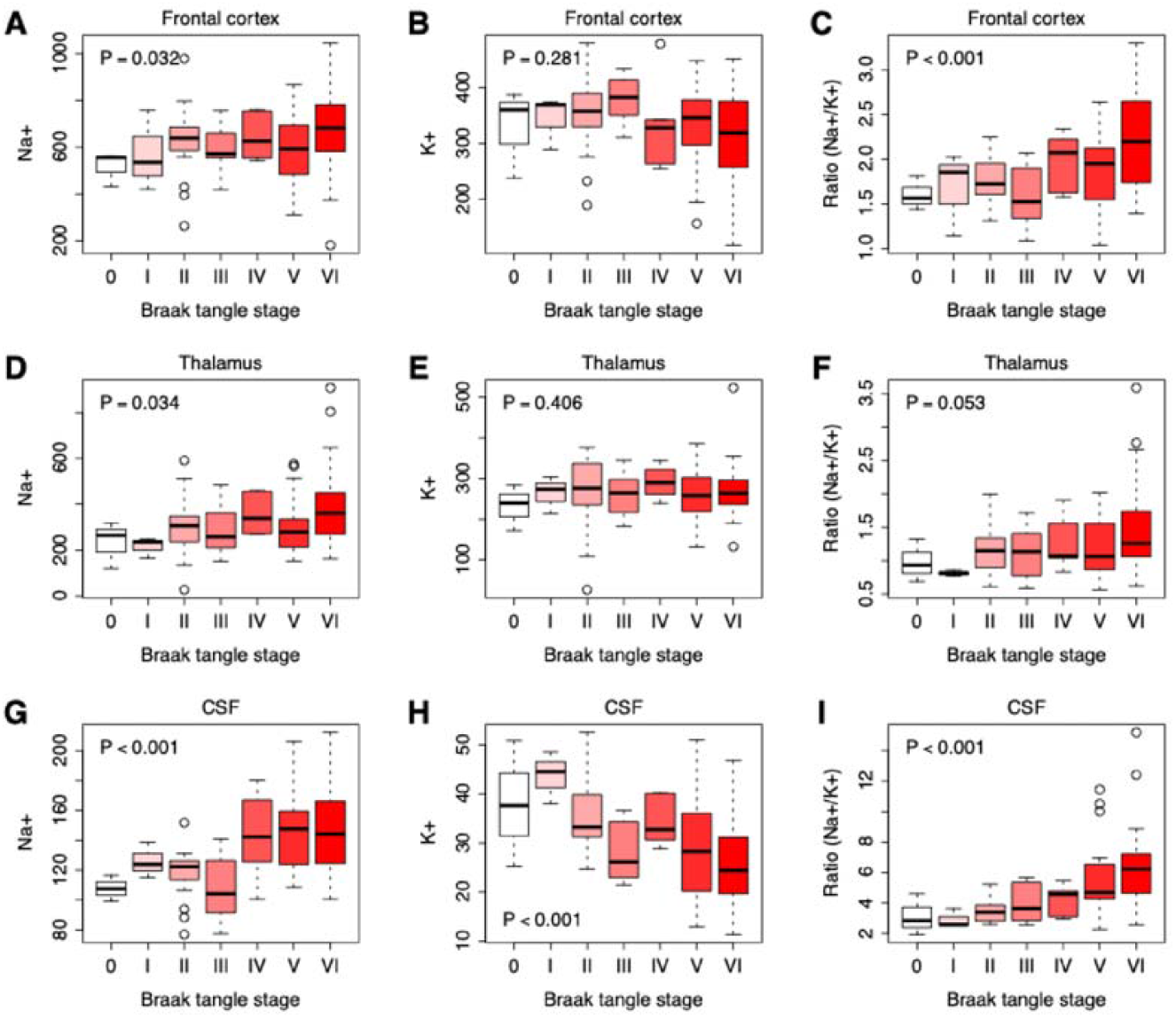
Na^+^, K^+^, or Na^+^/K^+^ ratio in the frontal cortex, thalamus, and CSF at Braak stages 0-VI. **(A-C)** The Na^+^ (A), K^+^ (B), or Na^+^/K^+^ ratio (C) in the frontal cortex at Braak tangle stage 1-VI. **(D-F)** The Na^+^ (D), K^+^ (E), or Na^+^/K^+^ ratio (F) in the thalamus at Braak tangle stage 1-VI. **(G-I)** The Na+ (G), K+ (H), or Na^+^/K^+^ ratio (I) in the CSF at Braak tangle stage 1-VI. Braak stage 0: N = 3; Braak stage I: N = 3; Braak stage II: N = 19; Braak stage III: N = 8; Braak stage VI: N = 6, Braak stage V: N = 27; Braak stage VI: N = 31. The P-values were computed by multivariate linear regression of Na^+^, K^+^, and Na^+^/K^+^ ratio on Braak stage, controlling for sex and age. Differences were considered statistically significant at *P*-values < 0.05.

## 3. Discussion

Epidemiological studies, clinical trials, and experiments in animal models provide compelling evidence that high dietary salt intake is causally linked to impaired cognitive function in AD and vascular dementia [36-39]. Nevertheless, data on whether Na^+^ accumulation is elevated in brain tissues and CSF in AD patients remains scarce. In this study, we measured [Na^+^] and [K^+^] in the prefrontal cortex, thalamus, and CSF in postmortem samples from human subjects. Several exciting observations were made: **1)** [Na^+^] was increased in the CSF and thalamus of AD subjects; **2)** the Na^+^/K^+^ ratio was significantly higher in brain tissues (prefrontal cortex and thalamus) and CSF of AD subjects; and **3)** the Na^+^/K^+^ ratio was significantly correlated with Braak stage. The higher the Na^+^/K^+^ ratio, the higher the Braak Stage (i.e., severity of AD tau pathology) in brain tissues and CSF, suggesting a strong association with disease progression.

A key finding of this study is elevated [Na^+^] in the CSF and thalamus of AD subjects, suggesting a potential disruption in Na^+^ homeostasis. These results align with the findings of Kerl et al. [40], who reported higher Na^+^ signal intensity in the CSF of early-stage AD patients compared with controls. This is despite methodological differences (^23^Na magnetic resonance imaging) of measurement and the living patients compared to our study of post-mortem brain tissue. Interestingly, our CSF-tissue correlation analysis revealed no significant relationship between CSF [Na^+^] and Na^+^ content in the thalamus or frontal cortex in both AD and control subjects. This suggests that brain tissue Na^+^ homeostasis is not simply a result of passive diffusion from the CSF, but rather reflects active regulation because passive diffusion would have resulted in a significant correlation between CSF-tissue ion levels. These findings also support previous studies indicating that dysfunction of Na^+^ transporters—responsible for maintaining physiological [Na^+^] in the CSF—may contribute to AD pathology [5,19,40,41]. One limitation of this study was the unavailability of plasma/serum samples from the same subjects, which is a challenge in post-mortem tissue studies because the bio sampling would need to be close to the time of death. Thus, we were unable to determine whether elevated plasma [Na^+^] contributes to increased CSF or brain tissue [Na^+^]. However, previous studies [42] have reported no significant correlation between CSF and plasma Na^+^ levels, reinforcing the concept of compartmentalized Na^+^ regulation in the brain. Another limitation of this study is that we were unable to evaluate the impact of potential confounders, such as post-mortem interval, time to freezing, number of freeze–thaw cycles, storage duration, or the use of antihypertensives or corticosteroids, which may influence the study’s conclusions.

Our study identified an elevation of the Na^+^/K^+^ ratio in brain tissues (prefrontal cortex and thalamus) and CSF of AD subjects. Using the Na^+^/K^+^ ratio as a measure helps to avoid the potential confounding effects of fluid volume in tissues on Na^+^ and K^+^ content. For example, fluctuations in brain fluid volume caused by edema can impact measurements of [Na^+^], which might not accurately reflect the actual Na^+^ content in the tissue (mEq/mg dry weight). The Na+/K+ ratio could therefore be a better marker for evaluating Na+ and K+ homeostasis.

The elevated Na^+^/K^+^ ratio in AD brain tissues aligns with the previous finding that AD is associated with Na^+^/K^+^-ATPase impairment [5,43]. Because of its electrogenic properties, Na^+^/K^+^-ATPase, which actively transports three Na^+^ ions out of the cell for every two K+ ions pumped in, plays a critical role in maintaining neuronal resting membrane potential and excitability. Dysfunction in this transporter can lead to intracellular Na^+^ accumulation and excessive K^+^ efflux, resulting in elevated intracellular [Na^+^] and reduced [K^+^] [44-46]. In other words, such dysfunction might give rise to an increased Na^+^/K^+^ ratio. Additionally, Na^+^/K^+^-ATPase in the choroid plexus is known to influence CSF Na^+^ levels in salt-sensitive hypertension models, likely due to increased Na^+^/K^+^-ATPase activity [47,48]. However, whether Na^+^/K^+^-ATPase activity is altered in the choroid plexus in AD remains unclear. The observed increase in CSF [Na^+^] and Na^+^/K^+^ ratio supports a hypothesis that elevated Na^+^/K^+^-ATPase activity in the choroid plexus may be present in AD pathology, though further research is needed to confirm this and to try and determine whether it might be a mediator or consequence of the disease.

Our study demonstrated that both [Na^+^] and the Na^+^/K^+^ ratio in CSF and tissues (thalamus and frontal cortex) of the brain are positively correlated with the severity of Braak tangle stage, suggesting the potential clinical relevance of these parameters in AD progression. Previous studies have reported similar correlations for Na^+^ levels and Braak stage [40,49]. For instance, Graham et al. measured Na^+^ in postmortem brain tissue of AD patients using ICP-MS and found a positive correlation between Na^+^ and Braak tangle stage [26]. Likewise, Mohamed et al. [49] used ^23^Na MRI and demonstrated a positive correlation between Na^+^ signal intensity in the hippocampus and the visual atrophy level of the medial temporal lobe, implying that Na^+^ increases with AD progression. Kerl et al. also observed a correlation between Na^+^ signal intensity and atrophy levels in the hippocampus and amygdala using ^23^Na MRI [40]. However, none of these previous studies assessed the Na^+^/K^+^ ratio.

Our study highlights the potential importance of this ratio in evaluating AD progression, providing a novel perspective on electrolyte balance in neurodegeneration. An interesting observation in our study is that higher Na^+^/K^+^ ratios were detected in both the prefrontal cortex and thalamus in AD, whereas no meaningful Na^+^ elevation was observed. This suggests that the Na^+^/K^+^ ratio may be a more sensitive indicator than absolute Na^+^ or K^+^, as it reflects changes in either or both ions (i.e., higher Na^+^ and/or lower K^+^ levels). Recent studies have suggested that the Na^+^/K^+^ ratio is a more sensitive marker compared with [Na^+^] or [K^+^] alone in predicting blood pressure outcomes and incident hypertension [27-29]. Given our findings, we propose that the Na^+^/K^+^ ratio could also serve as a valuable biomarker for assessing AD severity.

## 4. Materials and Methods

### 4.1. Human subjects

Brain prefrontal lobe cortex and thalamus tissues, and subject-matched CSF samples, from a total of 97 human subjects with (n = 67) and without (n = 30) AD were collected by post-mortem autopsy, frozen and stored at -80ºC before being shipped on dry ice. Tissues were taken from subjects selected according to their clinical and pathological diagnosis, categorizing them as either AD or normal, according to the detailed protocols of updated National Institute on Aging/Reagan Institute of the Alzheimer Association Consensus Recommendations for the Postmortem Diagnosis of AD (also known as the NIA-Reagan Criteria) that are widely accepted internationally recognized consensus criteria [50]. The tissues were obtained from the Human Tissue Authority-licensed South West Dementia Brain Bank (https://www.bristol.ac.uk/translational-health-sciences/research/neurosciences/research/south-west-dementia-brain-bank/), accessed on March 24^th^, 2020. Based on their clinical diagnosis from the NIA-Reagan Criteria [50], subjects were first categorized into two groups: control (age 50 years and older, with no or age-associated neuropathological abnormalities, Tau-related Braak tangle Stages 0-III) and AD (Braak tangle stage of IV-VI). Data for this study were collected between 1989 and 2015. Clinical and pathological reports, which include data on patient history, diagnosis, and de-identified personal information, were available for retrospective analysis. The South West Dementia Brain Bank is a Research Tissue Bank with ethical approval from the South West– Central Bristol Research Ethics Committee. The Research Integrity Offices at the University of Nevada, Reno, and the University of Bristol have determined that this project complies with human research protection oversight by the Institutional Review Board.

### 4.2. Flame photometry measurement of [Na^+^] and [K^+^] in the CSF and brain tissues

For the extraction of Na^+^ and K^+^ from the prefrontal lobe cortex and thalamus, 100 mg of wet tissues was heated first at 95°C for 48 hours and then at 105°C for an additional 48 hours, as previously described [51]. Dried tissue samples were digested in 2 mL HCl (1 mol/L) for 1 week. Then, each sample solution was centrifuged at 4°C at 10,000 x g and the supernatant was collected and diluted 10 times with distilled water. Na^+^ and K^+^ in the CSF and supernatants extracted from tissues were measured with a flame photometer (BWB BIO 943 Flame Photometer) [42] according to the manufacturer’s specifications, and their concentrations were determined by reference to a five-point calibration curve with inclusion of quality controls. [Na^+^] and [K^+^] in tissues were expressed as the standardized unit, μEq/g dry weight, representing the amount of Na^+^ or K^+^ per gram of dried brain tissue.

### 4.3. Statistical analysis

Data are presented as means ± SEM and were analyzed and plotted using Prism 10 software (GraphPad, La Jolla, CA, USA). Age demographics were assessed with an unpaired Student’s *t*-test. Differences in race demographics were assessed using Fischer’s test. The significance of differences between replicate means was assessed using two-way analysis of variance (ANOVA) with Fisher’s LSD post hoc analysis (multiple groups), as appropriate. Multivariate linear regressions were performed using the “lm” function in the R programming platform. All statistical tests were two-tailed, and differences were considered statistically significant at *P*-values < 0.05.

## 5. Conclusions and Perspectives

Our study provides compelling evidence that Na^+^ dysregulation and an elevated Na^+^/K^+^ ratio are associated with AD and its progression. Importantly, the Na^+^/K^+^ ratio, which we believe to be a more meaningful marker of Na^+^ and K^+^ homeostasis, was consistently higher in AD subjects across all examined regions, and positively correlated with Braak tangle stage, a widely used neuropathological marker of disease severity. These findings highlight the potential clinical relevance of ionic imbalances in AD and suggest that the Na^+^/K^+^ ratio, perhaps measured in CSF, may serve as a novel biomarker for disease progression.

These findings open several avenues for future research. First, further investigation into the mechanisms underlying Na^+^/K^+^-ATPase dysfunction, in conjunction with the measurement of levels of Na^+^ and K^+^ in AD is warranted to assess if impairment of this key ion transporter may explain the intracellular Na^+^ accumulation and K^+^ depletion. Additionally, studies examining whether CSF [Na^+^] and Na^+^/K^+^ ratio could serve as early diagnostic markers for AD patients are needed, perhaps alongside existing CSF Amyloid-Tao-Neurodegeneration biomarker approaches used already in clinical cohorts, but also in longitudinal cohorts tracking disease progression [52-54].

Recent evidence, including the study by Sasahara *et al*. suggests that amyloid β (Aβ) assemblies directly bind to and inhibit Na^+^/K^+^-ATPase, disrupting ionic gradients and contributing to AD pathogenesis [55-57]. Thus, another critical area for future research is the potential therapeutic implications of the modulation of Na^+^/K^+^ ratio in AD given that Na^+^/K^+^-ATPase is essential for maintaining neuronal membrane potential and cellular homeostasis. Targeting Na^+^ and K^+^ transporters such as Na^+^/K^+^-ATPase may offer new therapeutic strategies, aimed at enhancing its activity, rather than inhibiting it, which may prove beneficial in AD. Future therapeutic development might include small molecules that allosterically enhance ATPase activity, agents that prevent Aβ binding to the pump, or interventions that increase expression of specific isoforms of the ATPase that are more resistant to Aβ-induced inhibition.

Collectively, the results of our study highlight the importance of ionic balance in AD pathology and underscore the potential of the Na^+^/K^+^ ratio as a biomarker of disease severity. Understanding the mechanisms that drive Na^+^/K^+^ ratio dysregulation could provide valuable insights into AD progression and open new therapeutic avenues aimed at restoring ionic homeostasis in the aging brain.

## Author Contributions

Y.M.: Writing—review and editing, Writing—original draft, Methodology, Data curation, Investigation. S.P.: Writing—review and editing, Investigation, Methodology. T.Z.: Writing—review and editing, Formal Analysis. P.G.K.: Writing—review and editing, Conceptualization. Y.F.E.: Writing—review and editing, Conceptualization, Project administration, Funding acquisition. All authors have read and agreed to the published version of the manuscript.

## Institutional Review Board Statement

The Human Tissue Authority licensed South West Dementia Brain Bank, University of Bristol, which performed the post-mortem analysis, with tissue bank ethics approval from the South West–Central Bristol Research Ethics Committee (981105-1, approval date: 2016-11-03). The Research Integrity Offices at the University of Nevada, Reno, and the University of Bristol have determined that this project complies with human research protection oversight by the Institutional Review Board.

## Acknowledgments

We especially thank Dr. Laura Palmer for tissue processing and facilitating access to clinical data, and the South West Dementia Brain Bank (SWDBB), their donors and donor’s families for donating brain tissue for this study. Tissue for this study was provided with support from the BDR programme, jointly funded by Alzheimer’s Research UK and Alzheimer’s Society, and BRACE

