## Supplemental Figures for "Elevated Na^+^/K^+^ Ratio in Alzheimer’s Disease: A Potential Biomarker for Braak Stage"

University of Rochester Medical Center

601 Elmwood Avenue,

Rochester, NY 14642, USA

**Figure S1**

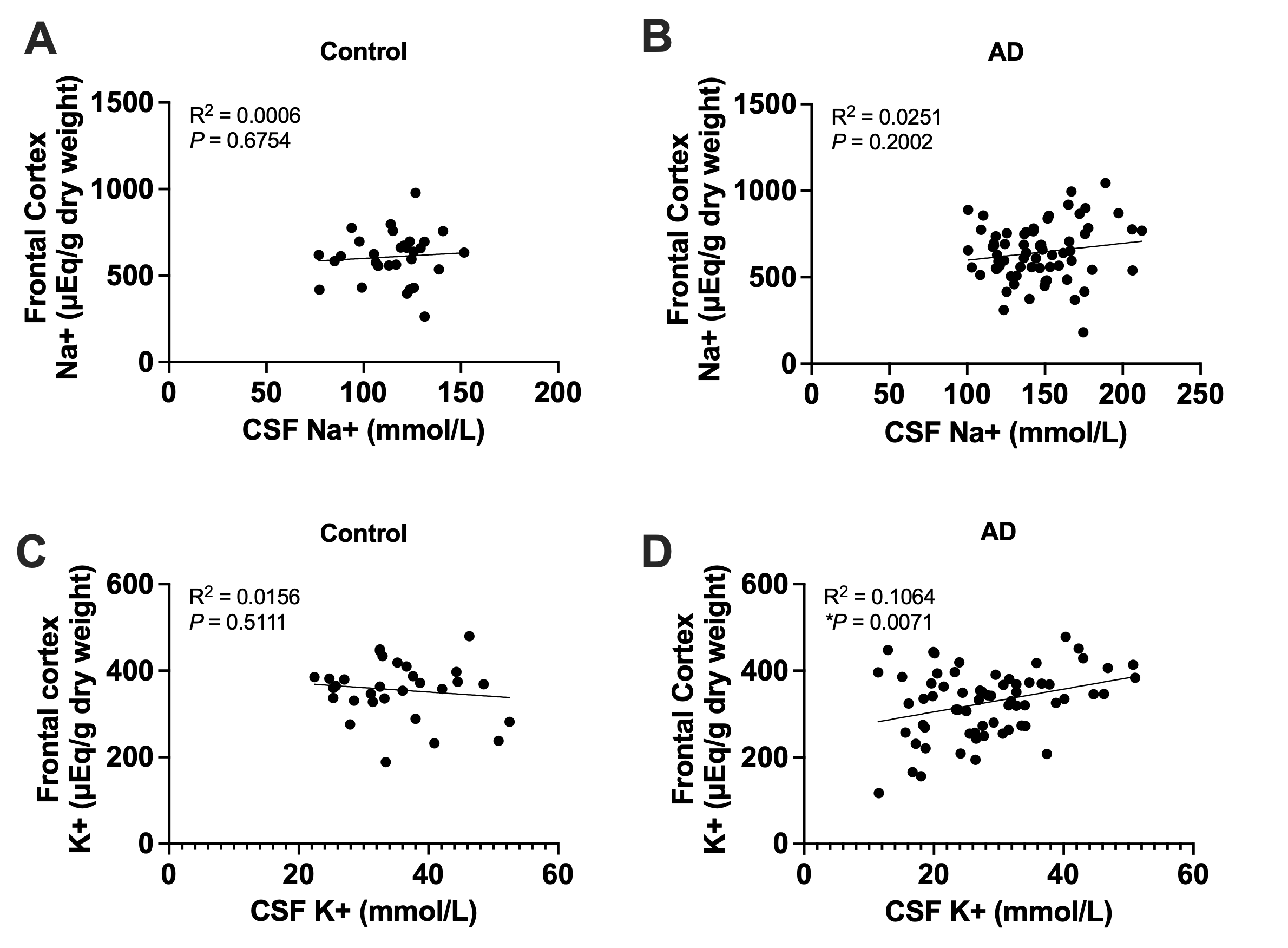

**Figure S1. Correlations between frontal cortex and CSF Na⁺, K⁺, and Na⁺/K⁺ ratio in control and AD subjects.** Linear regression analyses were performed to assess correlations between ion concentrations in the frontal cortex and cerebrospinal fluid (CSF) in control and Alzheimer’s disease (AD) subjects. (A,B) Frontal cortex versus CSF [Na⁺] in control (A; P = 0.6754, R² = 0.0006) and AD (B; P = 0.2002, R² = 0.0251) groups. (C, D) Frontal cortex versus CSF [K⁺] in control (C; P = 0.5111, R² = 0.0156) and AD (D; P = 0.0071, R² = 0.1064) groups.

**Figure S2**

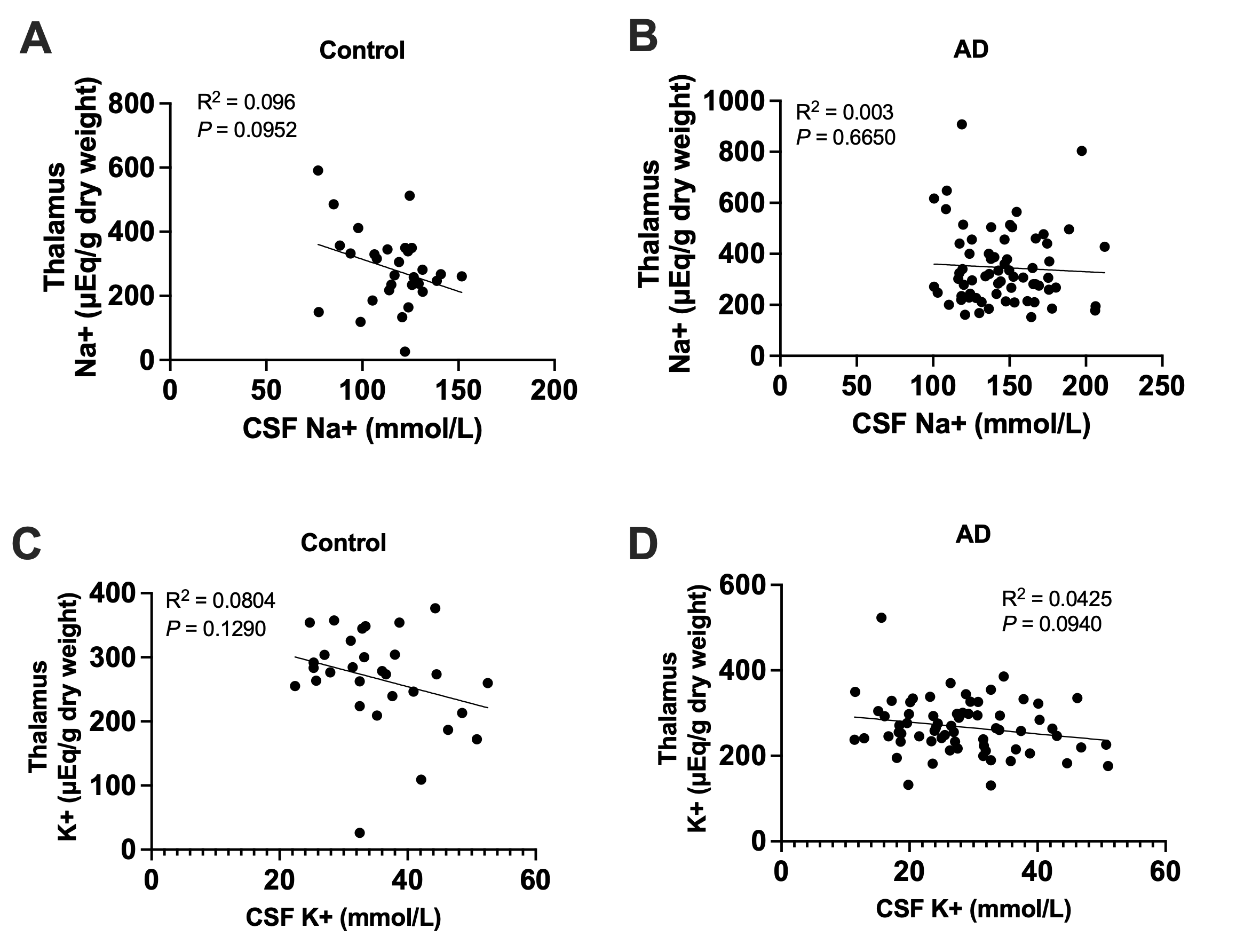

**Figure S2. Correlations between thalamic and CSF Na⁺, K⁺, and Na⁺/K⁺ ratio in control and AD subjects.** Linear regression analyses were performed to assess correlations between ion concentrations in the thalamus and cerebrospinal fluid (CSF) in control and Alzheimer’s disease (AD) subjects. (A, B) Thalamic versus CSF [Na⁺] in control (A; P = 0.0952, R² = 0.096) and AD (B; P = 0.6650, R² = 0.003) groups. (C, D) Thalamic versus CSF [K⁺] in control (C; P = 0.1290, R² = 0.0804) and AD (D; P = 0.0940, R² = 0.0425) groups.
